# A spinoparabrachial circuit defined by Tacr1 expression drives pain

**DOI:** 10.1101/2020.07.15.205484

**Authors:** Arnab Barik, Anupama Sathyamurthy, James Thompson, Mathew Seltzer, Ariel Levine, Alexander Chesler

## Abstract

Painful stimuli evoke a mixture of sensations, negative emotions and behaviors. These myriad effects are thought to be produced by parallel ascending circuits working in combination. Here we describe a pathway from spinal cord to brain for ongoing pain. Activation of a defined subset of spinal projection neurons expressing Tacr1 evokes a full repertoire of somatotopically-directed coping behaviors in the absence of noxious input. These cells project to a tiny cluster of Tacr1-positive neurons in the superior lateral parabrachial nucleus (PBN-SL) that themselves are responsive to sustained but not acute noxious stimuli. Activation of these PBN-SL^Tacr1^ neurons alone does not trigger pain responses but instead serves to dramatically heighten nocifensive behaviors and suppress itch. Remarkably, mice with silenced PBN-SL^Tacr1^ neurons ignore long-lasting noxious stimuli. These data reveal a spinoparabrachial pathway that plays a key role in the sensation of ongoing pain.

## Introduction

Sensory information about real and potential bodily damage reaches the brain through spinal projection neurons. Spinal projection neurons are sparse and heterogenous in their gene expression and their anatomical location (Koch et al., 2018; Peirs and Seal, 2016; Todd, 2010). For example, one group resides in the most superficial part of the dorsal horn (Lamina I) and is thought to receive input from peripheral nociceptors, itch neurons and thermoreceptors (Bester et al., 2000; Wercberger et al., n.d.; Wercberger and Basbaum, 2019). A second anatomically distinct set of projection neurons is localized to the deep dorsal horn (Laminae II-VI) where they integrate noxious and low-threshold sensory inputs (Wall, 1967; Wercberger and Basbaum, 2019; Willis, 1983). Both groups of spinal projection neurons target multiple sites in the brainstem and thalamus, with each site proposed to have unique roles in driving different aspects of sensation, learning and behavior (Huang et al., 2019; Liu, 1986; Todd, 2010). The prevailing view is that these parallel outputs combine to elicit the full pain experience. If true, then direct activation of the correct ensemble of projection neurons should produce behaviors indistinguishable from those evoked by actual noxious stimuli. By contrast, activation or silencing individual central targets might be expected to selectively affect particular aspects of pain and potentially reveal substrates that encode specific attributes of this complex state. Here we demonstrate that a spinoparabrachial circuit defined by the expression of the neuropeptide receptor Tacr1 is both necessary and sufficient for the heightened behavioral responses observed during ongoing noxious stimulation. Direct activation of the spinal Tacr1 neurons causes striking behaviors that closely match pain responses. Interestingly, activation of their parabrachial Tacr1 targets demonstrates that this branch of the ascending pathway is responsible for controlling the magnitude of the behavioral response and appears required for affective aspects of pain sensation.

## Results and discussion

We set out to define how activation of select ascending pathways contributes to pain sensation by probing a specific type of projection neuron (spinal^Tacr1^) previously reported to be needed for normal nociceptive behaviors (Mantyh et al., 1997). To probe the role of spinal^Tacr1^ neurons in pain behavior we developed a strategy to directly stimulate a small circumscribed group of these neurons using chemogenetics. Tacr1^Cre^ mice were transduced with an AAV encoding the excitatory DREADD receptor hM3Dq fused to an mCherry reporter (Barik et al., 2018; Krashes et al., 2011) on the right side of superficial lumbar dorsal horn (Figure 1A). As expected for the known expression of Tacr1 in spinal projection neurons (Todd, 2010), immunohistochemical analysis confirmed expression was restricted to population of sparse neurons in Lamina I and a second group of large neurons in Laminae III-V on the side ipsilateral to the injection location (L4; Figure 1B). Importantly, administration of the hM3Dq ligand clozapine-N-oxide (CNO) resulted in widespread Fos expression on the injected side (Figure 1B) validating our stimulation paradigm.

**Figure 1:**
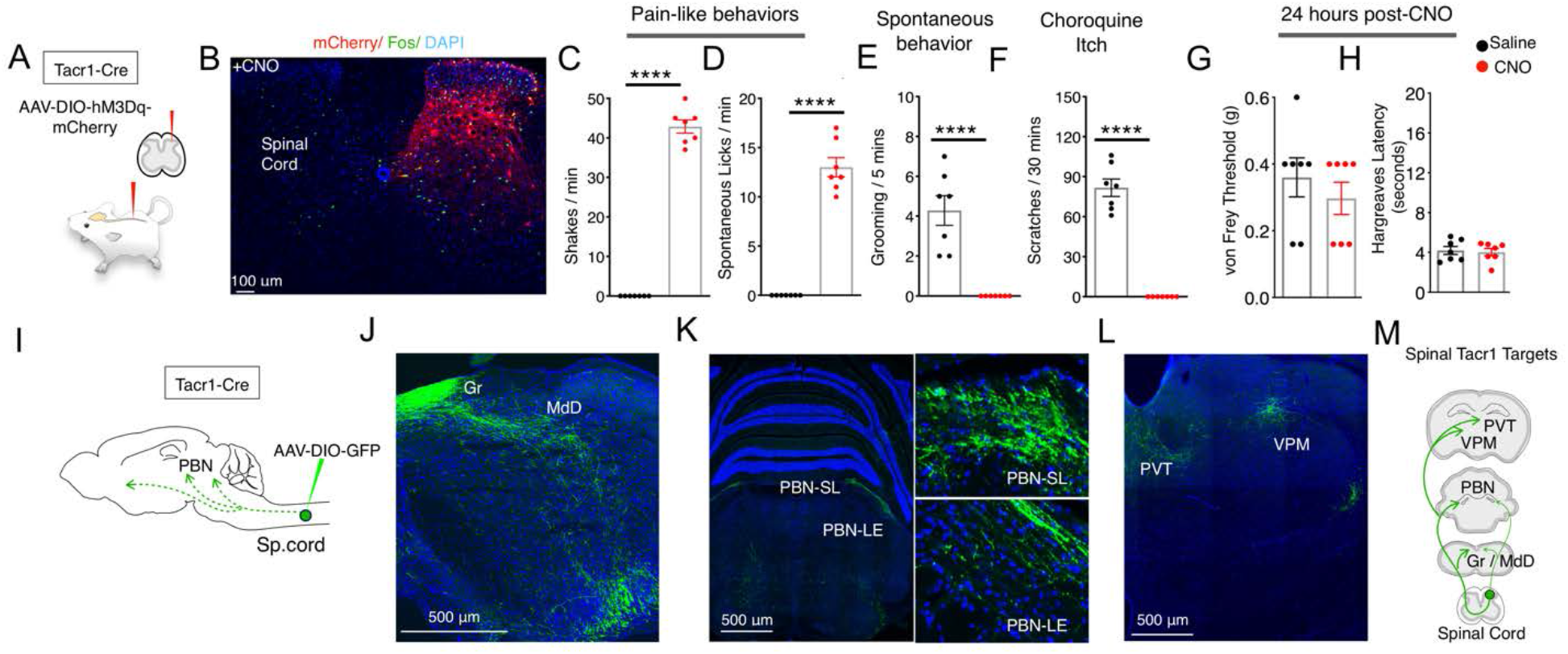
Activation of Tacr1+ projection neurons causes fictive pain. (A) A cartoon depicting the strategy to stimulate Tacr1-positive spinal neurons. An AAV viral vector encoding a Cre-dependent stimulating DREADD receptor (AAV-DIO-hM3Dq-mCherry) was injected into the lumbar spinal cord of Tacr1-Cre mice to transduce a localized subset of Tacr1-neurons. (B) A confocal image of a coronal section of lumbar spinal cord from an injected Tacr1-Cre mouse shows that the hM3Dq expression (red) is limited to neurons in the ipsilateral superficial and deep dorsal horn. CNO administration results in induction of the activity-dependent gene Fos (green). Scale = 100 μm (C-H) Bar graphs showing quantified CNO induced behavioral responses; saline control, black; CNO, red; n=7 mice/group; **** p ≤ 0.0001, unpaired two-tailed t test. CNO induced typical pain responses: ipsilateral hindpaw shaking (C) and licking (D) suppressed normal grooming (E) and itch responses to chloroquine injection in a different somatotopic site (F). As expected, behavioral changes were fully reversible after 24 h (G-H): shown are threshold responses to punctate touch (von Frey in grams; G) and latencies to respond to radiant heat (Hargreaves test; H). (I) A cartoon depicting the strategy to label the ascending projections of a small circumscribed group of Tacr1+ projection neurons. An AAV viral vector encoding a Cre-dependent GFP marker (AAV-DIO-GFP) was injected into the L4 region of the lumbar spinal cord of Tacr1-Cre mice. (J-L) Coronal sections showing major targets of Spinal^Tacr1^ neurons. In the brainstem, dense GFP labeled projections were detected in (J) the gracile nucleus (Gr) reticular formation (MdD) and (K) two regions of the parabrachial nucleus (PBN-EL and PBN-SL). GFP-positive Spinal^Tacr1^ projections are also observed in midline brain regions: the paraventricular thalamus (PVT) and ventral posterior medial thalamus (VPM). (M) A cartoon summarizing the ascending pathway of spinal Tacr1 expressing projection neurons. Green = GFP; Blue = DAPI. Scales = 500 μm.

Each side of a lumbar spinal cord segment receives sensory inputs from the ipsilateral hindlimb (Basbaum et al., 2009). Within minutes of CNO application, animals with spinal^Tacr1^ neurons expressing hM3Dq began spontaneously lifting, shaking and licking the hindlimb ipsilateral to the AAV injection (Figure 1C and D). These responses were highly reminiscent of those caused by extremely noxious and persistent stimuli such as chemical irritants (Kwan et al., 2006), lasted for over an hour and suppressed normal homecage behaviors (e.g. grooming; Figure 1E). Remarkably, animals on CNO became so focused on the DREADD-induced “fictive” pain associated with the ipsilateral hindlimb that they ignored other highly salient stimuli. In particular, they were completely unresponsive to chloroquine, a potent puritogen (Han et al., 2013; Mishra and Hoon, 2013) that normally causes robust and long lasting scratching (Figure 1F). Notably, chloroquine was applied to the nape of the neck, a completely different body area. Thus, in addition to directing nocifensive behavior to a somatotopically-conscribed area, the activation of a small group of lumbar spinal^Tacr1^ neurons produced the well-known phenomenon of itch suppression by pain (Ikoma et al., 2006). As expected, the effects of CNO were fully reversible and did not cause any long-lasting sensitization (Figure 1G and H).

We were surprised that artificial stimulation of the spinal^Tacr1^ neurons so accurately phenocopied behaviors caused by genuine noxious stimulation since various types of dorsal horn projection neurons are considered necessary for pain behavior (Koch et al., 2018; Todd, 2010). Interestingly, a recent study showed that ablating spinal neurons expressing *Tac1*, the gene encoding the neuropeptide Substance P, significantly impaired licking responses to sustained noxious stimuli (Huang et al., 2019). In the spinal cord, *Tac1* is broadly expressed both in projection neurons and interneurons (Gutierrez-Mecinas et al., 2017; Huang et al., 2019), however, our in situ hybridization data (Figure 1 - supplementary Figure 1A) revealed cells with high levels of Tacr1 transcripts in the superficial dorsal horn co-express Tac1. We reasoned that activation of spinal^Tac1^ neurons might produce similar behavioral effects as above (Figure 1). As predicted, chemogenetic stimulation of lumbar spinal^Tac1^ neurons (Figure 1 - supplementary Figure 1B, C and D) resulted in robust nocifensive behaviours directed to the ipsilateral hindlimb while at the same time suppressed grooming (Figure 1, Supplementary Figure 1E) and itch responses (Figure 1 - supplementary Figure 1F). There were minor differences in the behaviors with stimulation of Tac1^Cre^ animals being more robust perhaps in part due to activation of Tac1 expressing sensory neurons in the dorsal root ganglion (Figure 1 - supplementary Figure 1G). Together, these data show that a key population of dorsal horn neurons can be defined by the co-expression Tac1/Tacr1. Importantly, our results demonstrate that activity of the spinal cells expressing these genes carries the information needed to elicit a pain sensation localized to a specific body area.

Pain has many dimensions including the location, quality, intensity and duration. It also affects physiology, emotion, attention and memory (Bushnell et al., 2013). Output from the spinal cord dorsal horn targets multiple brain areas that work in concert to generate this complex state (Basbaum et al., 2009; Todd, 2010). We transduced lumbar spinal^Tacr1^ neurons with an AAV-DIO-GFP to visualize their supraspinal projections (Figure 1I). Consistent with their ability to coordinate a full-blown pain response, anterograde tracing revealed the GFP^+^ axons in many regions of the brain (Figure 1J through M). We focused on the PBN because of its importance in pain behavior (Barik et al., 2018; Barik and Chesler, 2020; Chiang et al., 2020; Huang et al., 2019). Outputs from two PBN areas, the lateral external and dorsal region are known to have diverse impacts on pain responses, threat learning and escape behaviors (Barik and Chesler, 2020; Campos et al., 2018; Chiang et al., 2020; Han et al., 2015). Although GFP^+^ axons targeted both these regions, the highest density spinal^Tacr1^ projections were at the very top of superior lateral PBN (PBN-SL; Figure 1K), a small and anatomically distinct area with unknown function. As expected, chemogenetic activation of spinal^Tacr1^ (Figure 2A) and spinal^Tac1^ (Figure 1, Supplementary Figure 1H) neurons induced robust Fos expression in the PBN-SL.

**Figure 2:**
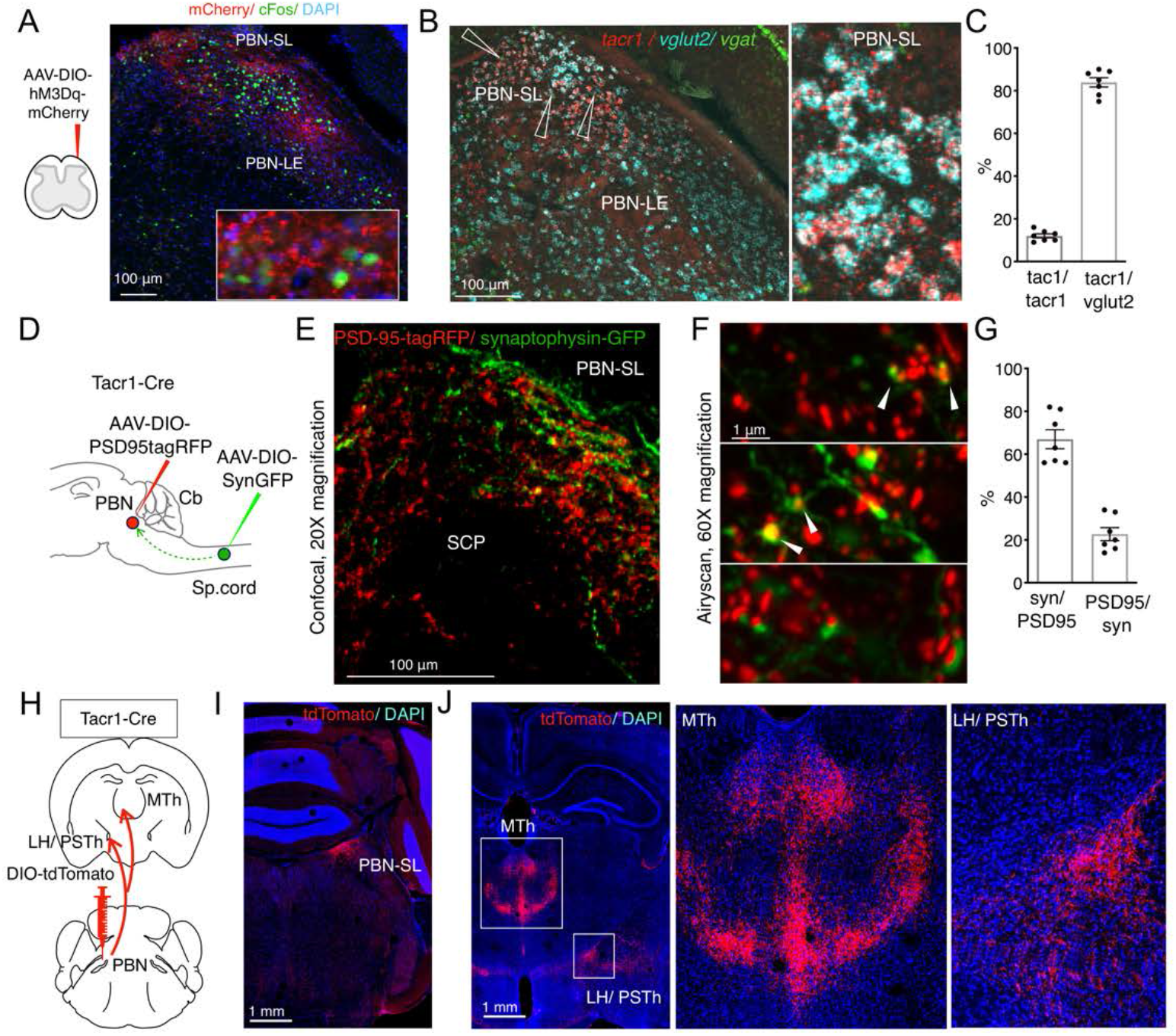
Anatomical organization of a Tacr1 defined spinoparabrachial circuit. (A) Spinal^Tacr1^ neurons were transduced to express hM3Dq-mCherry (cartoon, left); typical confocal image of a coronal section of the PBN showing mCherry positive Spinal^Tacr1^ projections (right panel, red) in two regions of the PBN (PBN-EL and PBN-SL). Application of CNO resulted in Fos induction throughout the PBN (green). The nuclear stain DAPI (blue) highlights the overall anatomy of the region. Scale = 100 μm. (B) Multichannel in situ hybridization shows Tacr1 transcript (red) is localized to the PBN-SL. Co-staining for the glutamate vesicular transporter Vglut2 (cyan) and GABA vesicular transporter Vgat (green) reveals most Tacr1-positive PBN-SL neurons are glutamatergic and hence excitatory. (C) Quantification of the percentage of Tacr1 positive cells co-expressing Tac1 (left plot) and Vglut2 (right plot). (D) A cartoon depicting the strategy used to label spinal^Tacr1^ presynaptic specializations and PBN-SL^Tacr1^ postsynaptic specializations in the same animal. Cre-dependent viral vectors were injected in the lumbar spinal cord (AAV-DIO-SynGFP to label presynaptic termini) and PBN (AAV-PSD95tagRFP to label postsynaptic densities) in Tacr1^Cre^ mice. (E) Example confocal image of a coronal section from the PBN showing a low magnification view of the organization of Spinal^Tacr1^ presynaptic specializations (green) and PBN-SL^Tacr1^ postsynaptic specializations (red). Scale = 100 μm.(F) Super-resolution imaging (Airyscan) of sections from 3 different mice showing close apposition of SynGFP and PSD95tagRFP puncta indicative of synaptic connections. Scale = 1 μm. (G) Quantification of number of SynGFP puncta with PSD95tagRFP puncta in close apposition (left graph) and vice versa (right graph) demonstrate that the majority of spinal projection neurons target PBN-SL^Tacr1^ neurons; n= 7 sections from n=3 mice. (H) A cartoon depicting the viral strategy for anterograde tracing of PBN-SL^Tacr1^ neuron projections using injection of AAV-DIO-tdTomato into the PBN of Tacr1^Cre^ mice. (I-J) Confocal images of coronal sections showing tdTomato labeling (red) of cell bodies of PBN-SL^Tacr1^ neurons (I) and their projections (J). Dense projections were found in two major brain regions: the medial thalamus (MTh) and a region encompassing part of the lateral hypothalamus (LH) and the parasubthalamic nucleus (PSTh). Scale = 1 mm.

Which cells in the PBN-SL receive the major input from the spinal dorsal horn and how do they contribute to pain sensation? To answer these questions, we sought molecular markers that could be used to genetically manipulate PBN-SL neurons so we could study their anatomy and function. Since both the neuropeptide (Tac1) and one of its receptors (Tacr1) were expressed by the spinal projection neurons (Figure 1 - supplementary Figure 1A) we predicted that tachykinin signaling molecules might mark this sensory circuit. Indeed, multiplex fluorescent in-situ hybridization revealed Tacr1 to be highly expressed in a tight cluster of excitatory cells in the PBN-SL (Figure 2B, C and Figure 2 - supplementary Figure 1). The expression of Tacr1 in both the spinal and PBN-SL neurons meant their anatomical connectivity could be probed by differentially transducing each region with AAVs in the same Tacr1^Cre^ animal (Figure 2D). We labeled the presynaptic specializations of spinal^Tacr1^ axons with Synaptophysin-GFP and the postsynaptic specializations of the PBN-SL^Tacr1^ dendrites and soma using PSD95-tagRFP (Figure 2E). Near super-resolution imaging of single Z planes from stained tissue sections from the PBN-SL showed very close apposition of red and green varicosities indicative of functional synapses (Figure 2F, G).

Despite its compact structure, the PBN appears central to many different sensory pathways with the various anatomic regions selectively targeting distinct higher centers and controlling different aspects of visceral sensation (Bernard et al., 1994), thermoregulation (Yahiro et al., 2017; Yang et al., n.d.), itch (Campos et al., 2018; Mu et al., 2017) and pain (Bernard and Besson, 1990; Campos et al., 2018; Menendez et al., 1996). Therefore, we next used anterograde tracing to determine the projections of PBN-SL^Tacr1^ neurons by targeting the PBN of Tacr1^Cre^ animals with an AAV-DIO-tdTomato (Figure 2H, I). Two forebrain regions were prominently innervated by PBN-SL^Tacr1^ neurons (Figure 2J): (1) the midline thalamus (MTh) and (2) the lateral hypothalamus/ parasubthalamic nucleus (LH/PSTh). Both these regions appear to be selective targets of PBN-SL excitatory projections since they have not been previously shown to be innervated by PBN neurons (Barik et al., 2018). Moreover, expression of synaptophysin-GFP in PBN-SL^Tacr1^ neurons provided evidence for presynaptic terminal specializations in both areas (Figure 2 - supplementary Figure 2).

Do the same neurons target both MTh and LH/PSTh? Or are there two distinct types of PBN-SL^Tacr1^ neurons with different higher brain targets and potential roles? To answer these questions, we adopted an intersectional strategy where we labeled PBN-SL^Tacr1^ neurons projecting to the LH/PSTh. Our approach used a Cre-dependent mCherry to label the cell bodies of Tacr1-expressing neurons in the PBN and a Flp-dependent synaptophysin-GFP (Sathyamurthy et al., 2020) to label their synaptic termini (Figure 2 - supplementary Figure 3; see methods for details). We confirmed that the PBN-SL neurons expressed both labels (Figure 2 - supplementary Figure 3A) meaning that PBN-SL^Tacr1^ neurons project to the LH/PSTh, validating our approach. Importantly, Synaptophysin-GFP puncta were also abundant in the MTh (Figure 2 - supplementary Figure 3B) demonstrating collateralization of these cells. Together, these anatomical experiments reveal two previously unappreciated targets of PBN-SL^Tacr1^ neurons in the forebrain. Therefore, we anticipated that these neurons may play a specialized role related to pain and next set out to record their responses as well as determine their function.

**Figure 3:**
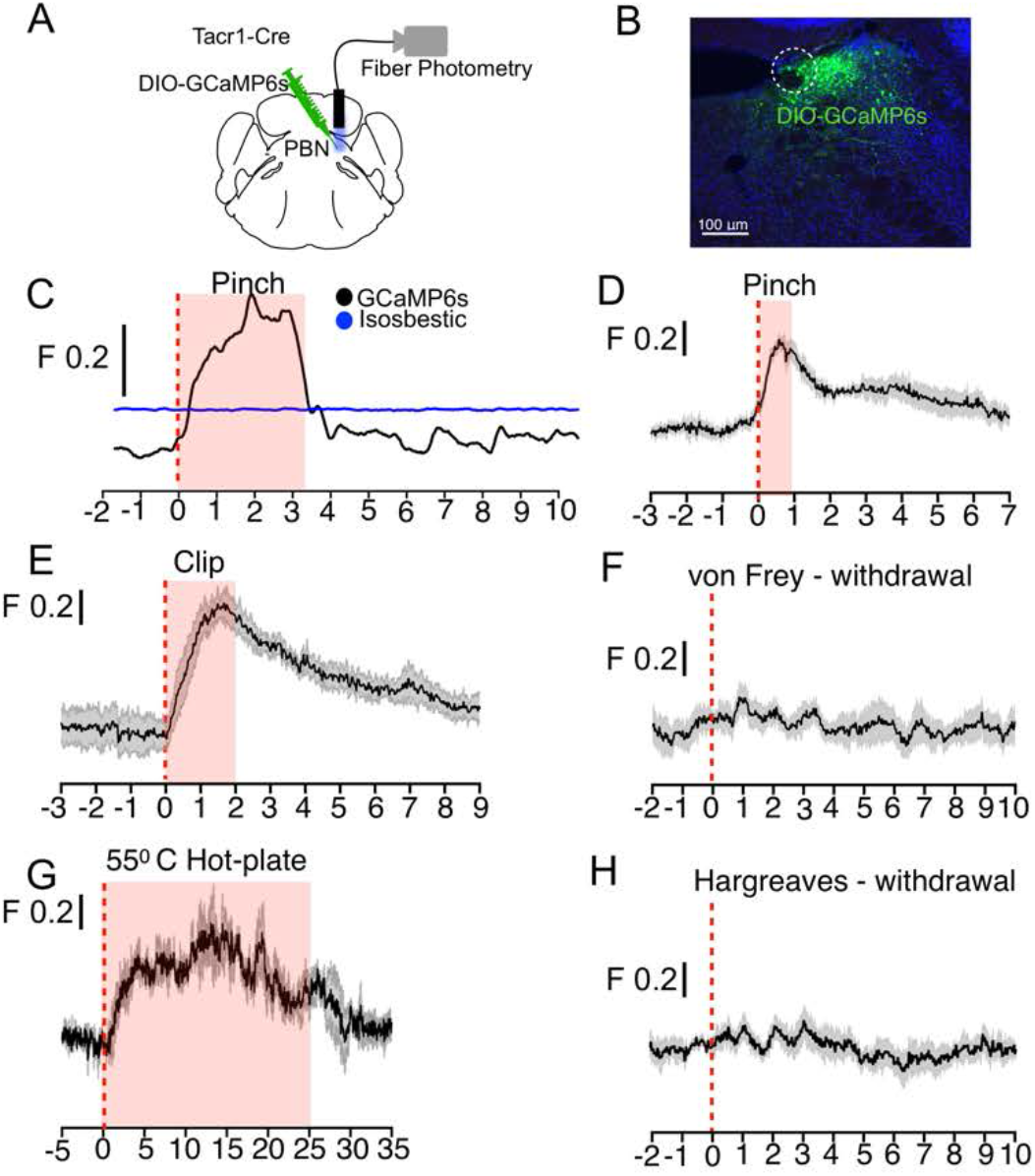
PBN-SL^Tacr1^ neurons respond to sustained noxious stimulation. (A) Cartoon depicting the approach for recording population activity in PBN-SL^Tacr1^ neurons: a Cre-dependent viral vector encoding the genetically encoded calcium indicator GCaMP6f was injected into the PBN of Tacr1^Cre^ mice and an optical fiber was placed over the injection site. Fiber photometry recording was used to monitor population calcium responses to different somatosensory stimuli. (B) After recording, posthoc staining and confocal imaging of PBN sections confirmed GCaMP6f expression (green) and fiber placement (dotted line). (C) An example photometry trace from a lightly anesthetized mouse showing a robust, time locked calcium response. The change in fluorescence/total fluorescence is shown (black line; F) to tail pinch with blunt forceps (stimulus onset at 0 seconds; dotted line). Note the response lasts the duration of the stimulation (shaded region) before returning to baseline. As a control for movement artifacts, 405 fluorescence (Blue trace; Isosbestic) was also monitored and showed no changes. (D-H) Average population calcium responses in awake mice to sustained (D, E and G) and acute (F, H) noxious stimuli. Dark lines are means of responses from multiple animals aligned to the start of stimulation (time 0) with the standard error shown in light grey. X-axis shows time in seconds aligned to the start of stimulation; F indicates ΔF/F scaling (D) Pinching (hindpaw) with a blunt forcep (n= 6 mice), (E) Clip assay applied to hindpaw (n=5 mice). (F) von Frey stimulation (0.6g) aligned to paw withdrawal (n=4 mice) (G) 55 °C Hot plate test aligned to when mice are placed in a chamber and lasting 25 seconds (n=3 mice). (H) Radiant heat test (Hargeaves) aligned to paw withdrawal (n=7 mice).

Tacr1 expression marks a rather homogeneous group of neurons in the PBN. Fiber photometry provides a simple yet sensitive approach for measuring the functional responses of genetically defined and anatomically restricted neuronal populations (Gunaydin et al., 2014). Therefore to study how PBN-SL^Tacr1^ neurons are tuned to respond to sensory stimuli, we engineered mice expressing the calcium sensor GCaMP6f (Chen et al., 2013) in these neurons and implanted optical fibers to record population level activity (Figure 3A, B). Consistent with our anatomical studies (Figure 2), fiber-photometry recordings demonstrated that many types of noxious stimulus strongly activate PBN-SL^Tacr1^ neurons. An example recording (Figure 3C) highlights the typical type of long-lasting calcium transient that was elicited by a single pinch of the hindpaw of a lightly anesthetized mouse. Activation of these neurons was observed irrespective of the pinch location (Figure 3 - supplementary Figure 1). Next we recorded activity in non-anesthetized mice using two types of noxious but non-damaging sustained mechanical stimulation: pinch with blunt forceps (Figure 3D) and pinch with the clip assay (Figure 3E). Again, PBN-SL^Tacr1^ neurons exhibited robust population calcium responses to this type of stimulation (Figure 3D, E). By contrast, punctate mechanical stimuli did not evoke calcium responses in PBN-SL^Tacr1^ neurons (Figure 3F). This was surprising since we used stiff von Frey filaments (0.6g) that reliably cause nocifensive reflexes (Abdus-Saboor et al., 2019) but may indicate that PBN-SL^Tacr1^ neurons preferentially respond to sustained stimuli.

Painful mechanical, thermal and chemical stimuli are detected by separate nociceptive channels (Basbaum et al., 2009; Gatto et al., 2019; Le Pichon and Chesler, 2014). We next asked whether these pathways converged on PBN-SL^Tacr1^ neurons by recording evoked activity in this population to high temperatures and chemical irritants. In keeping with these neurons playing a general role in pain sensation they exhibited robust activity to hot-plate exposure (Figure 3G) and topical application of the pungent component of mustard oil, allyl isothiocyanate (AITC; Figure 3 - supplementary Figure 2A, B). Like pinch, both of these are long lasting painful stimuli that activate distinct sets of peripheral nociceptors (Basbaum et al., 2009). However, PBN-SL^Tacr1^ neurons were not activated by short-term noxious heat (Hargreaves test; Figure 3H). Interestingly, although AITC is known to cause allodynia and hyperalgesia (Albin et al., 2008), this type of skin irritation did not sensitize PBN-SL^Tacr1^ responses to acute touch or heat stimuli (Figure 3 - supplementary Figure 2C-F). In combination, the photometry data strongly support PBN-SL^Tacr1^ neurons as playing a major role in an ascending pathway from spinal dorsal horn that is activated by sustained noxious inputs.

We next set out to determine how this circuit influences pain behaviors by generating mice where PBN-SL^Tacr1^ neurons could be chemogenetically activated (Figure 4A, B). Although activation of PBN-SL^Tacr1^ neurons did not induce spontaneous pain responses, CNO delivery transformed behavior: mice became skittish, avoided handling and repeatedly attempted to escape from the test chambers. This general heightened state of arousal and anxiety was quantified using an open field arena as increased movement and avoidance of the center (Figure 4 - supplementary Figure 1A). Mice now withdrew from the slightest touch with thin von Frey filaments (0.07g; Figure 4, Supplementary Figure 1B), something not seen in saline controls. Moreover, in a sustained pinch assay using a clip applied to the paw, activation of PBN-SL^Tacr1^ neurons increased nocifensive behavior and almost doubled the time a mouse spent shaking the clipped paw (Figure 4C, D). Even more strikingly, behavioral changes persisted after the clip was removed. Although saline controls quickly returned to normal resting behaviors, after CNO the same animals continued to lick and investigate the hindpaw for many minutes after the pinch was terminated (Figure 4E). Equally robust effects of stimulating PBN-SL^Tacr1^ neurons were observed in responses to AITC, a chemical irritant which normally causes licking and attending to the affected area (Figure 4F). The time a CNO-treated mouse spent licking the injected site more than doubled. CNO injected mice also exhibited heightened escape responses when exposed to heat in a hotplate assay (Figure 4G). Therefore, this circuit plays a role in controlling the magnitude of behavioral responses to a range of noxious stimuli that activate different peripheral sensory neurons. Interestingly, the latency in a mouse’s response to heat, thought to involve simple spinal reflexes, was not influenced indicating that peripheral sensitization is not involved (Figure 4 – supplementary Figure 1C and D). Thus, activation of PBN-SL^Tacr1^ neurons induces a “hyper-vigilant” state resembling the effects of long-term noxious stimulation (e.g. inflammation).

**Figure 4:**
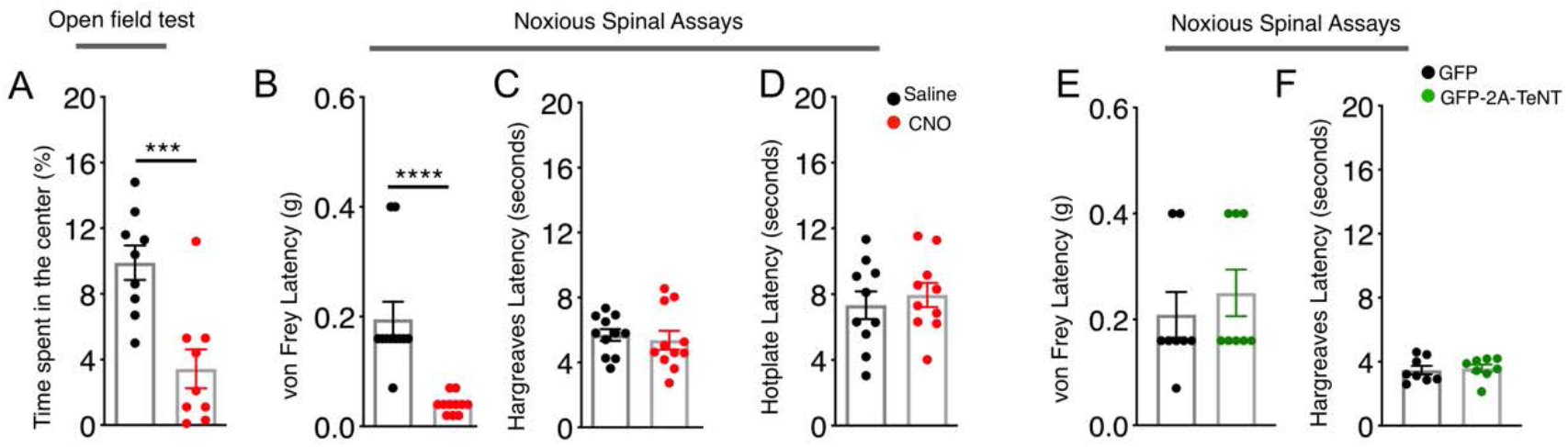
PBN-SL^Tacr1^ neurons are necessary and sufficient to drive pain-related behaviors. (A-H) Chemogenetic activation of PBN-SL^Tacr1^ neurons results in hyperalgesia. (A) A cartoon depicting the strategy for stimulation of PBN-SL^Tacr1^ neurons. The PBN of a Tacr1^Cre^ mouse was injected with a Cre-dependent viral vector encoding an excitatory DREADD receptor (AAV-DIO-hM3D1-mCherry) to allow chemogenetic activation. (B) Typical confocal images showing coronal sections of the PBN region from transduced Tacr1^Cre^ mice treated with saline (top image) or CNO (bottom image). The expression of mCherry in both animals is restricted to select neurons in the PBN-SL (red). CNO application but not saline injection results in induction of the activity-dependent gene Fos (green) validating the approach. Scale = 100 μm. (C-G) Chemogenetic stimulation of PBN-SL^Tacr1^ neurons heightens behavioral responses sustained by noxious mechanical (C-E), chemical (F) and thermal (G) stimulation (n=7 mice tested with saline, black points and then with CNO, red points on different days). Mice treated with CNO spend significantly more time shaking (C) and exhibit more licking events (D) directed to the hindpaw during the Clip assay. Notably, CNO treated mice continued to lick the hindpaw for several minutes after the clip was removed (E). CNO treatment also increased the number of times mice licked their paws after topical AITC treatment (F) and jumped to escape from a 55 °C hotplate (G). CNO treatment did not increase result in licking behaviors in the absence of noxious stimulation (H). (I-O) Silencing of PBN-SL^Tacr1^ neurons inhibits normal pain responses to sustained noxious stimuli. (I) A cartoon depicting the strategy for silence the output of PBN-SL^Tacr1^ neurons. Bilateral injection of a Cre-dependent viral vector encoding Tetanus Toxin Light C GFP fusion (AAV-DIO-GFP-2A-TeNT) in Tacr1^Cre^ mice was used to silence PBN-SL^Tacr1^ neurons; AAV-DIO-GFP served as a control. (J) Typical confocal image of a coronal section through the PBN region showing the restricted expression of GFP-2A-TeNT in the PBN-SL (green). Scale = 100 μm. (K-O) Silencing of PBN-SL^Tacr1^ neuronal output significantly dampened behavioral responses to sustained noxious mechanical (K-M), chemical (N) and thermal (O) stimulation (n=8 GFP, black points; n=7 TeNT-GFP, green points). In the Clip (sustained pinch assay, K-M), expression of GFP-2A-TeNT in PBN-SL^Tacr1^ neurons decreased the time spent shaking (K) and the number of licking events (L). Similarly, silencing PBN-SL^Tacr1^ neurons significantly decreased the number of times mice licked their paws after topical AITC treatment (N); since control animals did not make escape attempts from a 55 °C hot plate (O) no effect of neuronal silencing was observed in this assay. Unpaired two-tailed t test; ns > 0.05, *** p ≤ 0.001, **** p ≤ 0.0001.

Recent work from other groups has suggested that inhibiting PBN activity suppresses scratching (Campos et al., 2018; Mu et al., 2017), however our data show that stimulating the Tacr1 expressing spinal neurons strongly suppressed itch related behaviors. Therefore, we next investigated how activation of PBN-SL^Tacr1^ neurons affected behavioral responses to pruritogens. To induce itch, we injected chloroquine, a potent non-histaminergic puritogen that activates a select type of peripheral itch neuron and evokes vigorous scratching (Han et al., 2013). Injecting this compound into the nape of the neck of control mice reliably induced localized bouts of scratching starting a few minutes after injection and lasting for about 30 minutes. Remarkably, chemogenetic stimulation of the small group of PBN-SL^Tacr1^ neurons completely suppressed scratching (Figure 4 - supplementary Figure 2), mimicking the effects both of painful stimuli and the fictive pain induced by activating Tacr1 expressing spinal projection neurons. Therefore, the PBN-SL^Tacr1^ neurons serve as a central substrate that is sufficient to suppress itch.

Chemogenetic activation and functional recordings demonstrate that PBN-SL^Tacr1^ neurons respond to painful stimuli and can induce hypervigilance and increased behavioral responses to sustained types of pain. However, it is well known that the central pathways controlling pain sensation and responses involve many regions of the brain (Bushnell et al., 2013). Therefore we next used directed expression of tetanus toxin (TeNT-GFP) (Han et al., 2015; Li et al., 2019) to silence synaptic output and determine the role of this PBN-SL^Tacr1^ circuit in pain behaviors (Figure 4I, J). We again examined responses to different types of sustained stimuli that in control animals induce robust behaviors: the clip assay of extended pinch, and topical application of AITC. Remarkably, in both cases, silencing PBN-SL^Tacr1^ neurons almost completely eliminated shaking and licking normally induced by these stimuli (Figure 4K-O). Moreover, at a qualitative level, mice with silenced PBN-SL^Tacr1^ neurons effectively ignored these normally painful stimuli. Thus, this small and localized cluster of PBN-SL^Tacr1^ neurons appears to play a profound role in pain and is necessary for normal behavioral responses.

Together our results expose a circuit involving Tacr1 expressing neurons in the spinal cord and the PBN-SL that controls how mice respond to a wide range of persistently painful stimuli. Activating lumbar spinal^Tacr1^ projection neurons closely mimics pain by inducing pronounced, somatotopically directed behaviors. The PBN-SL^Tacr1^ neurons are one CNS-target of these projection neurons and are both necessary and sufficient to drive a subset of these pain-related behaviors. This small group of brainstem neurons are tuned to long lasting noxious input regardless of location and are selectively important for behaviors that only occur when pain is persistent. When activated, PBN-SL^Tacr1^ neurons dramatically potentiate these types of complex behavioral responses but not simple reflexes. Interestingly, when this group of neurons is stimulated in the absence of noxious peripheral input, mice become ‘jumpy’ and hypervigilant, now recoiling from the gentlest of touches. However, unlike spinal projection neurons, PBN-SL^Tacr1^ neurons do not themselves evoke pain responses or provide positional information but instead appear to drive affective aspects of pain. This is particularly apparent in the clip assay where mice with silenced PBN-SL^Tacr1^ neurons almost completely ignore this type of sustained, and normally painful, pinch. In the future, determining how other central targets of the spinal^Tacr1^ projection neurons control behavior should help reveal if pain responses are simply the sum of distinct parallel circuits or require interactions between distributed regions of the brain.

## Methods

### Mouse lines

Animal care and experimental procedures were performed in accordance with a protocol approved by the National Institute for Neurological Diseases and Stroke (NINDS) Animal Care and Use Committee. Tac1-Cre or Tac1-IRES2-Cre-D or B6;129S-Tac1tm1.1(cre)Hze/J (Stock number 021877) (Barik et al., 2018; Harris et al., 2014) mice were purchased from Jackson Laboratories. Tacr1-T2A-Cre-Neo (Tacr1-Cre) was kindly donated by Dr Hongkui Zheng, Allen Brain Institute. Genotyping for the mentioned strains was performed according to protocols provided by the Jackson Laboratories.

### Viral vectors and stereotaxic injections

Mice were administered 1 ml saline mixed with 25 mg/kg of ketoprofen 30 minutes prior to and for 2 days daily post-surgery. Mice were anesthetized with 2% isoflurane/oxygen prior and during the surgery. Craniotomy was performed at the marked point using a hand-held micro-drill (Roboz). A Hamilton syringe (5 or 10 ul) with a glass pulled needle was used to infuse 200 nl of viral particles (1:1 in saline) at 100 nl/min. The following coordinates were used to introduce the virus: PBN-SL-AP:-5, ML:±1.36; DV:3.30; MTh--AP:-1.72, ML:±0.2; DV:3.5. The stereotaxic surgeries to deliver AAVs in the lumbar spinal cord were performed as described before (Sathyamurthy et al., 2020). Vectors used and sources: AAV9-CAG-FLEX-tdTomato (UPenn; donated by Allen Institute); AAV9-CAG-FLEX-GFP(UPenn; donated by Allen Institute); AAV5-hSyn-DIO-mCherry (UNC); AAV5-hSyn-DIO-hM3Dq (UNC; donated by Brian Roth); AAV8-Flex-hSyn-Synaptophysin-YFP (MGH GDT Core); AAV9-Flex-hSyn-PSD95-TagRFP (this paper; donated by Mark Hoon); AAVDJ-CMV-eGFP-2A-TeNT (Stanford Viral Core GVVCAAV-71); AAVRetro-FLPO (Sathyamurthy et al., 2020)1-DIO(FRT)-Synaptophysin (Sathyamurthy et al., 2020). Post Hoc histological examination of each injected mouse was used to confirm viral-mediated expression was restricted to target nuclei.

### Behavioral assays

One experimenter carried out all the behavioral assays for the same cohort and was blinded to the treatments (J.T. or M.S.). Experiments throughout all the intraplantar and intraperitoneal administrations were performed by one experimenter (A.B.). All experiments were done in the same room, which was specifically designated for behavior and under red light. Mice were habituated in their home-cages for at least 30 minutes in the behavior room before experiments. For the Hargreaves test, mice were habituated in transparent Plexiglass chambers for 30 minutes. Clozapine-N-Oxide (1 mg/kg) dissolved in DMSO and diluted in saline was injected i.p. 60-90 minutes before behavioral experiments or histochemical analysis (Krashes et al., 2011).

The clip assay was performed as described in (Huang et al., 2019). For the AITC test, 10 % AITC (Sigma) in saline was injected intradermally in the paw as described in, (McNamara et al., 2007). All the experiments were videotaped with an over-head video camera or a panasonic digital camera and scored offline post-hoc using TopScan by CleverSys. The Hargreaves apparatus, and the programmable hotplate were purchased from IITC and used according to the manufacturer's instructions. Von Frey test was done by manually applying the following filaments: 0.008, 0.02, 0.04, 0.07, 0.16, 0.4, 0.6, 1, 1.4 g. Animals received 10 stimulations on the left paw. Inter trial interval was at least 15s. Open field was from CleverSys. Mice were always habituated in their respective chambers for 30 minutes prior experimentation.

### Fiber photometry

A three-channel fiber photometry system from Neurophotometrics was used to collect data. Light from two LEDs (470 nm and 405 nm) were bandpass filtered and passed through a 20 x Nikon Objective focussed on a fiber optic cable coupled to the cannula implanted on mouse PBN-SL. Fluorescence emission was collected through the same patch cord and filtered on a CMOS sensor. Data was acquired through Bonsai. Photometry data was analyzed using a custom MATLAB code provided by Neurophotometrics. To correct for photobleaching and heat mediated LED decay, the isosbestic signal was fit with a biexponential that was then linearly scaled to the calcium-dependent fluorescence signal F. The ΔF/F was calculated by dividing the signal by the scaled fit. The start and end of the stimuli (where applicable) were timestamped. Where possible, simultaneous video recording with a Microsoft webcam was performed.

### Immunostaining, multiplex in situ hybridization and confocal microscopy

Mice were anesthetized with isoflurane and perfused intracardially with PBS and 4% PFA (Electron Microscopy Sciences) consecutively for immunostaining experiments. Fresh brains were harvested for in situ hybridization experiments. For Fos experiments, brains were harvested 90 minutes after hotplate/ capsaicin/ ATP and 150 minutes after i.p. CNO administration or intraplantar formalin. Tissue sections were rinsed in PBS and incubated in a blocking buffer (5% goat serum; 0.1% Triton X-100; PBS) for 3 hours at room temperature. Sections were incubated in primary antibodies in the blocking buffer at 4°C overnight. Sections were rinsed 1-2 times with PBS and incubated for 2 hours in Alexa Fluor conjugated goat anti-rabbit/ rat/ chicken secondary antibodies (Thermo Fisher Scientific), secondary antibodies washed in PBS, and mounted in ProLong gold mounting media (Thermo Fisher Scientific) onto charged glass slides (Daigger Scientific). Multiplex ISH was done with a manual RNAscope assay (Advanced Cell Diagnostics). Probes were ordered from the ACD online catalog. Z stack images were collected using a 20X and 40X oil objective on a laser scanning confocal system (Olympus Fluoview FV1000) and processed using ImageJ/FIJI software (National Institutes of Health). The AiryScan images of synapses were acquired with 60X oil objective on a Zeiss LSM 880 AiryScan Confocal microscope.

### Statistical analyses

All statistical analyses were performed using GraphPad PRISM 7 software. ns > 0.05, * p ≤ 0.05, ** p ≤ 0.01, *** p ≤ 0.001, **** p ≤ 0.0001.

## Acknowledgements

We are indebted to Nicholas Ryba and Mark Hoon (NIDCR), as well as members of Chesler lab (NCCIH) for invaluable discussions and advice on the manuscript. We are grateful to Hongkui Zheng (Allen Brain Institute for providing the Tacr1-Cre mice) and Mark Hoon for the AAV-DIO-PSD95tagRFP. We thank Dr Carolyn Smith at the NINDS imaging core facility for help with acquiring AiryScan images. This work was supported by the Intramural Research Program of the NIH, National Center for Complementary and Integrative Health (ATC) and National Institute of Neurological Disorders and Stroke (ATC and AL).

---

**Figure 1 - supplementary Figure 1:**
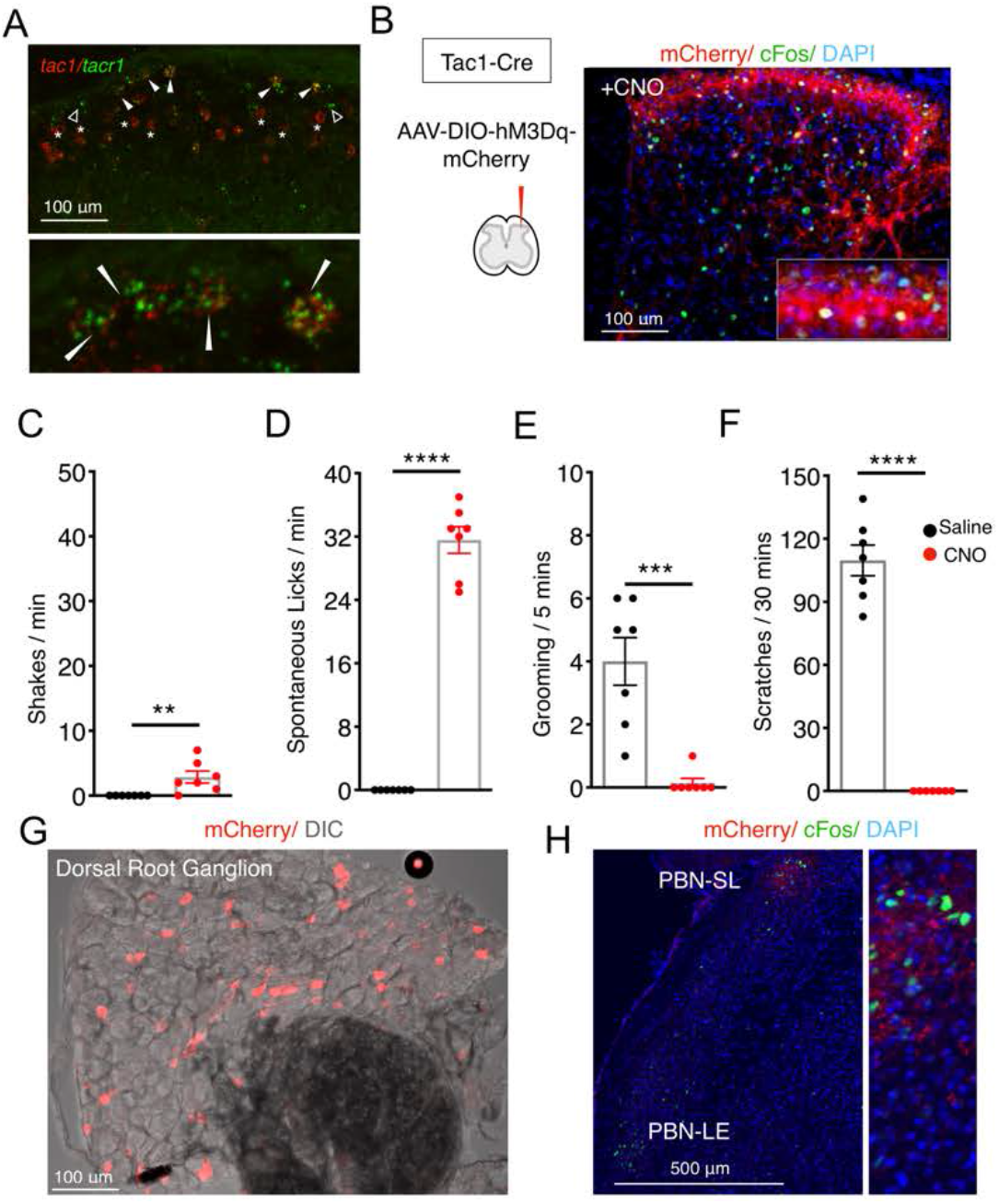
Tac1 and Tacr1 are co-expressed in nociceptive spinal projection neurons. (A) A confocal image of a coronal spinal cord section showing the detection of mRNA for Tac1 (red) and Tacr1 (green) using two color fluorescent in situ hybridization. Filled arrowheads indicate cells co-expressing both genes, open arrowheads are Tacr1-only cells. Note that there are many red cells that are not green (stars) below the superficial part of the dorsal horn (Lamina I) consistent with Tac1 also being expressed in excitatory interneurons. Lower panel is a higher magnification of the superficial dorsal horn region highlighting doubly positive cells (arrowheads). Scale = 100 μm. (B) An AAV viral strategy analogous to the one used (Figure 1) to activate Spinal^Tacr1^ neurons was used to express the excitatory DREADD (hM3Dq-mCherry) in lumbar Spinal^Tac1^ neurons. Confocal images of coronal spinal cord sections from infected mice after CNO administration and stained with DAPI (blue) revealed mCherry-positive neurons (red) and widespread induction of Fos (green). Inset shows a higher magnification view. Scale = 100 μm. (C-F) Stimulation of Spinal^Tac1^ neurons results in ‘fictive pain’ behaviors. Bar plots show the behaviors of mice (n=7) treated with saline (black) and CNO (red) show that hM3Dq mediated activation of Spinal^Tac1^ neurons results in shaking (C) and licking (D) of the ipsilateral hindlimb, suppression of normal grooming behaviors (E), and inhibition of responses to the pruritogen chloroquine applied to the nape of the neck (F). Unpaired two-tailed t test; **** p ≤ 0.0001. (G) Spinal injection of AAV-DIO-hM3Dq-mCherry in Tac1^Cre^ mice retrogradely labeled nociceptors in the dorsal root ganglia (red). Scale = 100 μm. (H) Spinal^Tac1^ neurons labeled with hM3Dq-mCherry project to the PBN-SL (red). CNO activation of these cells results in Fos induction (green) in this brain region. Scale = 500 μm.

**Figure 2 - supplementary Figure 1:**
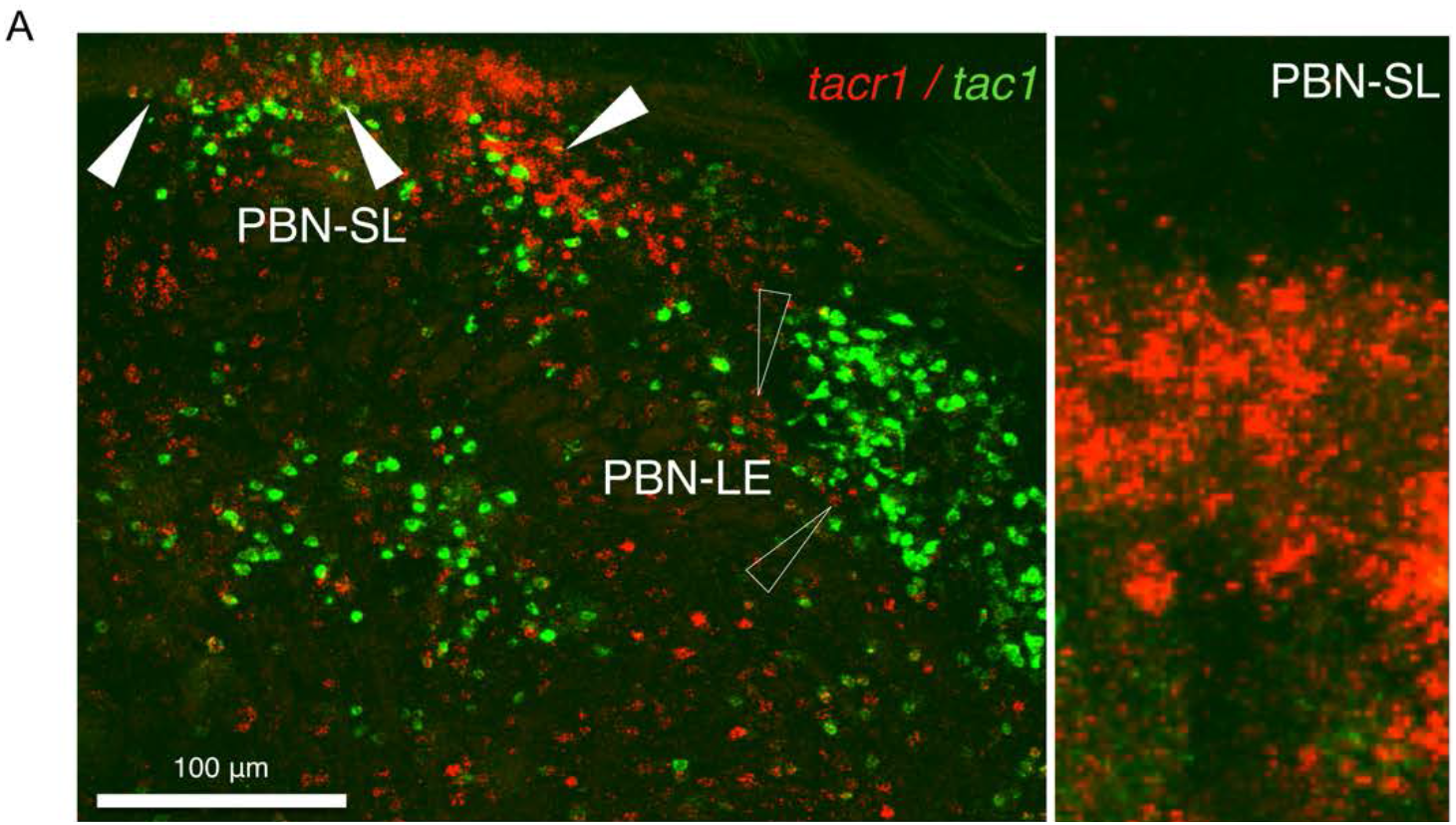
Tac1 and Tacr1 are differentially expressed in the PBN. Typical confocal image of two color in situ hybridization showing Tacr1 expression (red) is enriched in the PBN-SL and is largely non-overlapping with Tac1 (green). Right side is an enlargement of the PBN-SL showing densely clustered Tacr1 neurons. Scale = 100 μm.

**Figure 2 - supplementary Figure 2:**
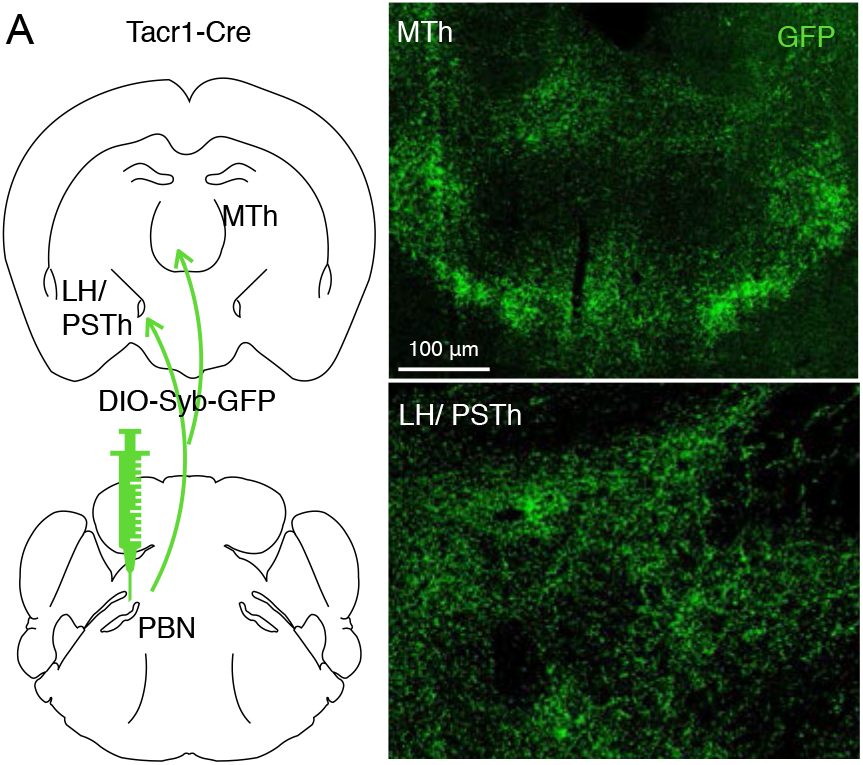
Viral-assisted visualization of PBN-SL^Tacr1^ projection terminals. The presynaptic terminals of PBN-SL^Tacr1^ neurons were labeled by transducing Tacr1^cre^ mice with a viral vector encoding a Cre-dependent synaptophysin-GFP fusion construct (AAV-DIO-SynGFP). Confocal imaging of coronal sections of the Medial Thalamus (MTh) and Lateral Hypothalamus/Parasubthalamic nucleus (LH/PSTh) reveal densities of GFP-positive puncta in both regions indicating that PBN-SL^Tacr1^ neurons make presynaptic terminals in both regions. Scale = 100 μm.

**Figure 2 - supplementary Figure 3:**
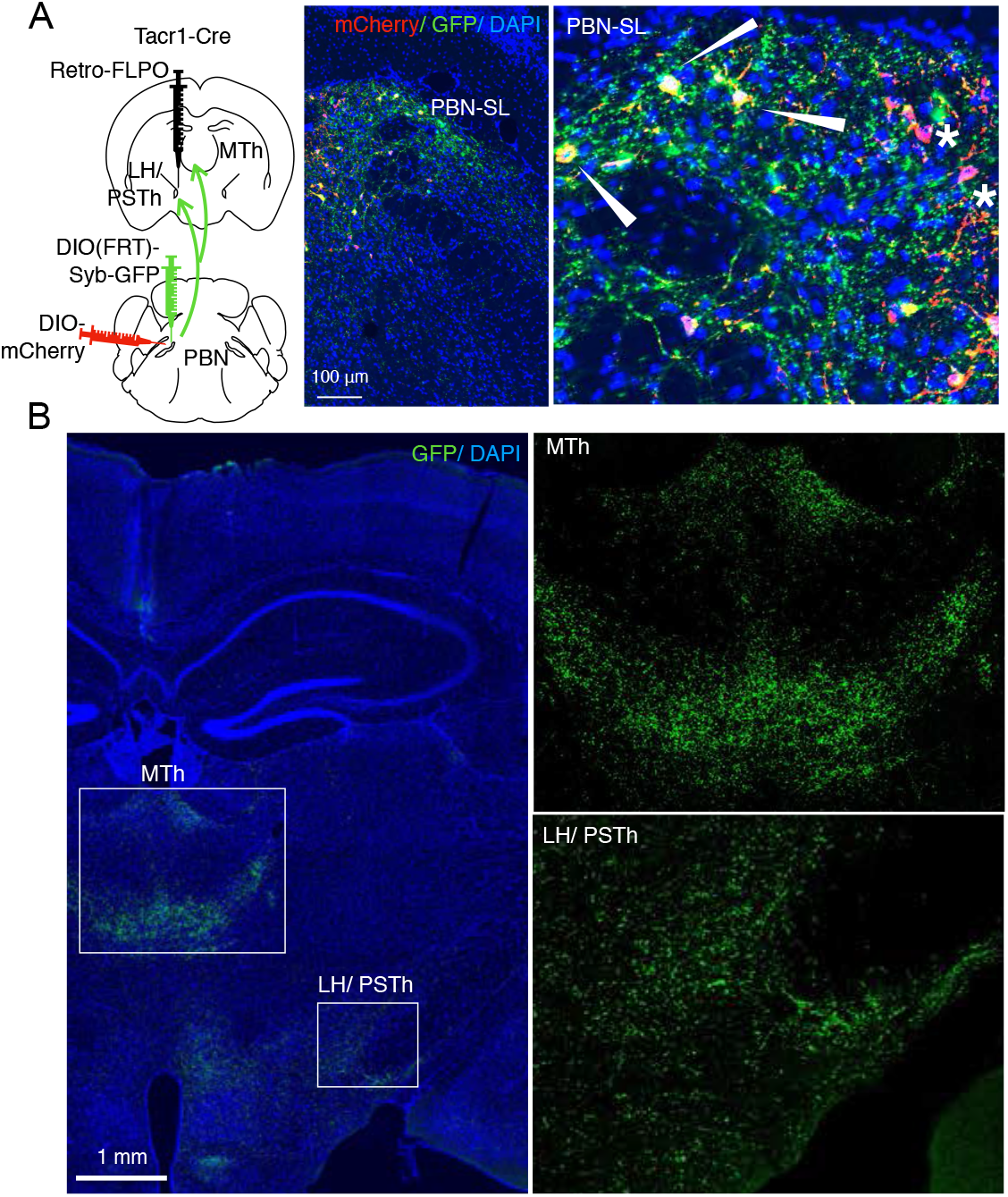
PBN-SL^Tacr1^ neurons collateralize and project to both MTh and LH/PSTh. (A) A complex intersectional genetic strategy was devised to label PBN-SL neurons that project to LH/PSTh and examine if they project to other brain areas (cartoon left side). The terminals of PBN neurons in the LH/PSTh were transduced with retrograde viral vector encoding Flp recombinase in a Tacr1^cre^ mouse. In the same animal, two viral additional vectors were co-injected into the PBN. The first encoded a Flp-dependent presynaptic marker (SynGFP) and the second encoded a Cre-dependent cellular marker (mCherry). Under these conditions, the cells bodies and axons of PBN neurons that project to LH/PSTh (and were transduced by the retrograde Flp-virus) are labeled in green and PBN-SL^Tacr1^ neurons are labeled in red. Confocal images of coronal sections of the PBN (right panels) show a high degree of overlap between red (mCherry) and green (SynGFP) confirming the PBN-SL^Tacr1^ neurons project to LH/PSTh (arrowheads). Note that not every cell is co-labelled, as expected when examining the intersecting expression of three different AAVs injected at two distinct sites (stars). Blue = DAPI stain; Scale = 100 μm. (B) PBN-SL neurons that project to LH/PSTh have collaterals that make presynaptic terminal specializations in the MTh. Confocal image of a coronal section of a mouse where PBN neurons that project to the LH/PSTh were labeled with SynGFP. As expected, many GFP positive puncta are seen in the LH/PSTh (boxed region, lower right). Notably, a high density of GFP positive puncta are also found in the MTh in the same section (boxed region, center). High magnifications of each boxed area are shown on the right side. Blue = DAPI stain; Scale = 1 mm.

**Figure 3 - supplementary Figure 1:**
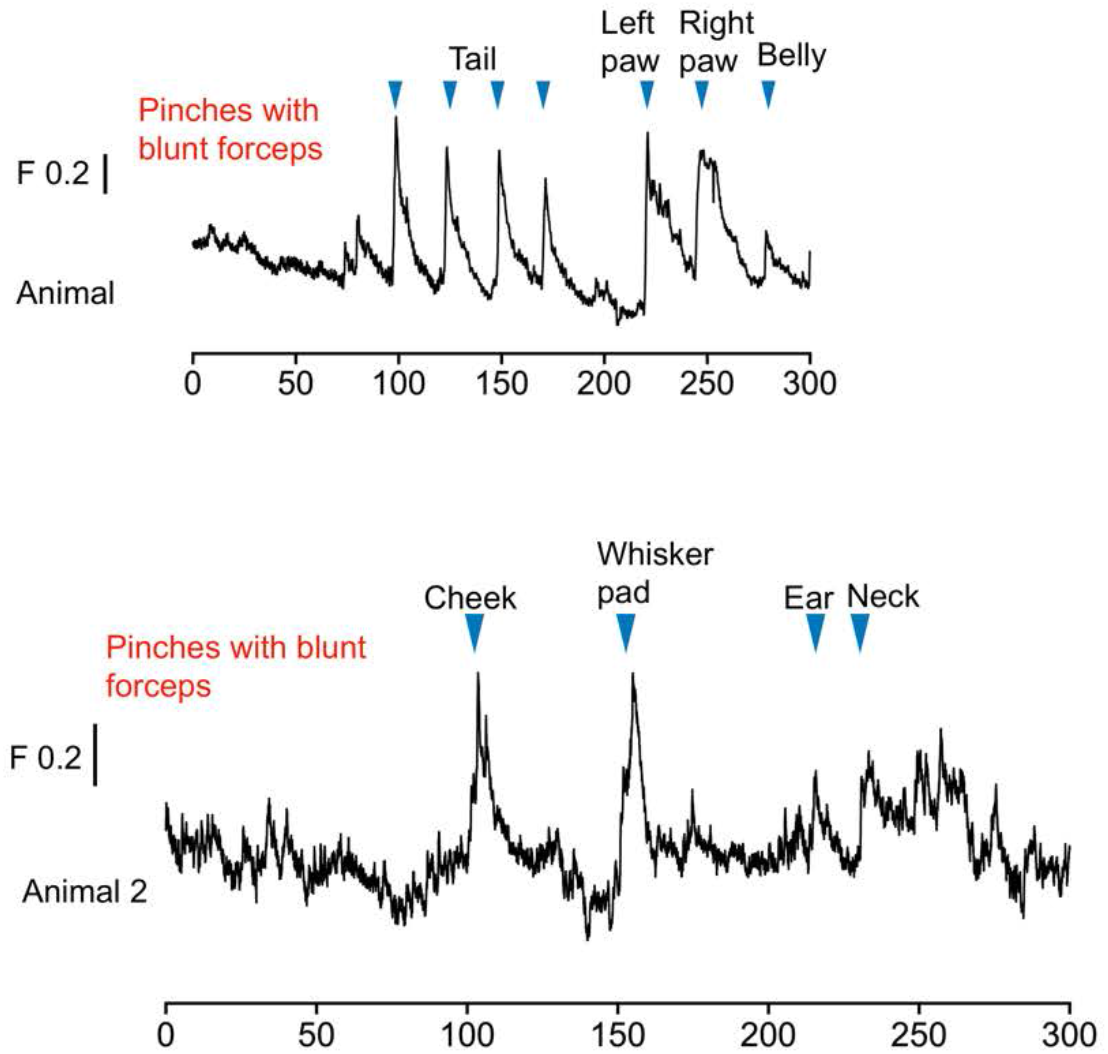
Population calcium responses in PBN-SL^Tacr1^ neurons to pinch are not somatotopically restricted. Example traces from two separate lightly anesthetized mice showing robust responses to a series of sustained pinches with blunt forceps (blue arrowheads) of different body parts (tail, left hindpaw, right hindpaw, belly, cheek, whisker pad, ear and neck). Time is in seconds, 0 is the start of the trail, change in fluorescence/total fluorescence is shown (black line; F).

**Figure 3 - supplementary Figure 2:**
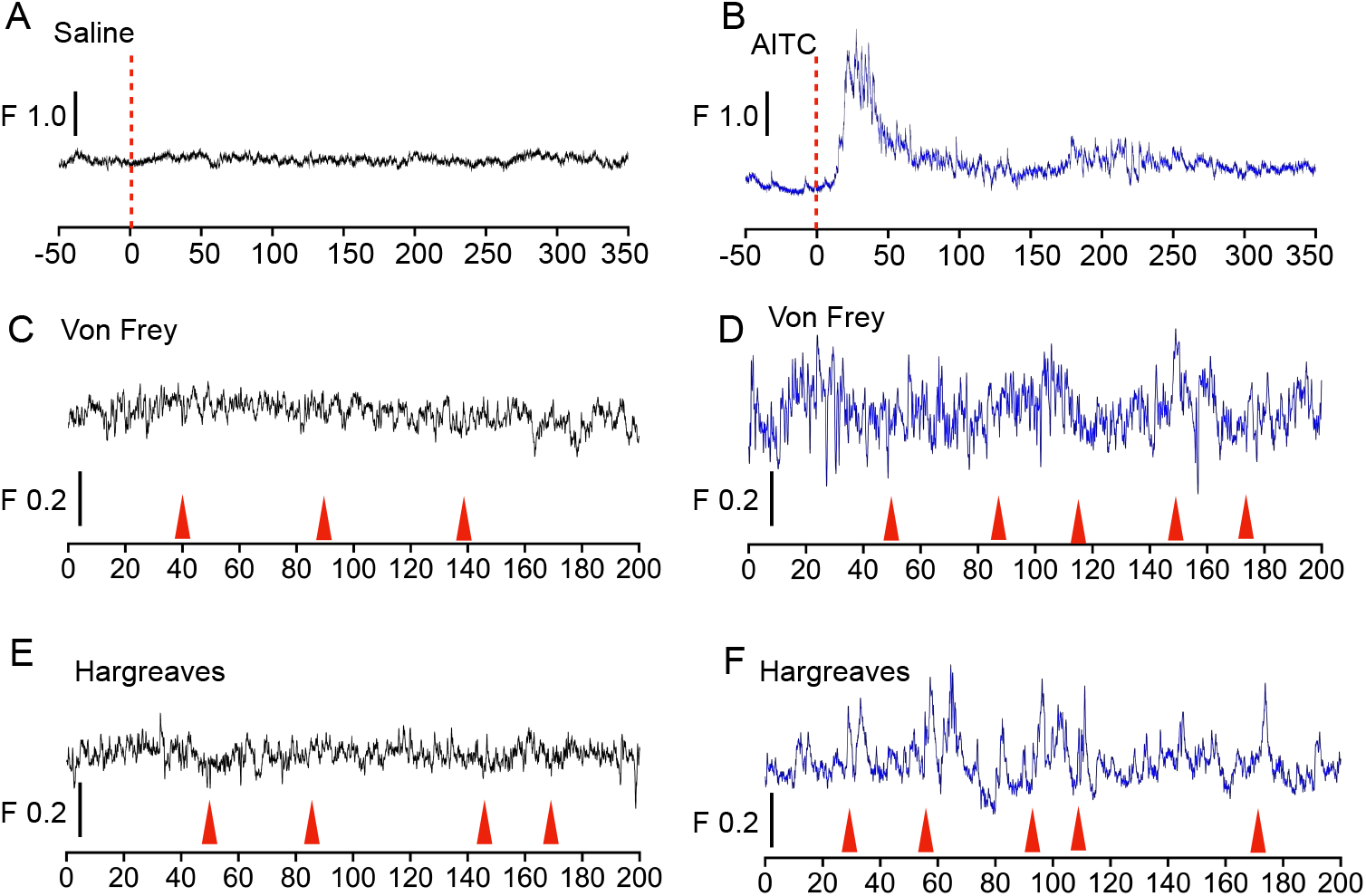
The chemical irritant allyl isothiocyanate (AITC) evokes large and sustained increases in calcium signaling in PBN-SL^Tacr1^ neurons. Example traces from an awake mouse to topical application of saline (A) and then (B) AITC on the hindpaw. AITC evokes a very robust response followed by long-lasting increases in spontaneous calcium transients. Note that despite the increased baseline activity and behavioral hyperalgesia after application, AITC treatment does not change the sensitivity of these neurons to acute noxious stimulation.Both acute noxious mechanical (C, D) or thermal (E, F) stimuli fail to activate PBN-SL^Tacr1^ neurons either in control or AITC treated conditions. Red arrows indicate behavioral responses (paw withdrawals). Time is in seconds, 0 is the start of the trail, change in fluorescence/total fluorescence is shown (black line; F).

**Figure 4 - supplementary Figure 1:**
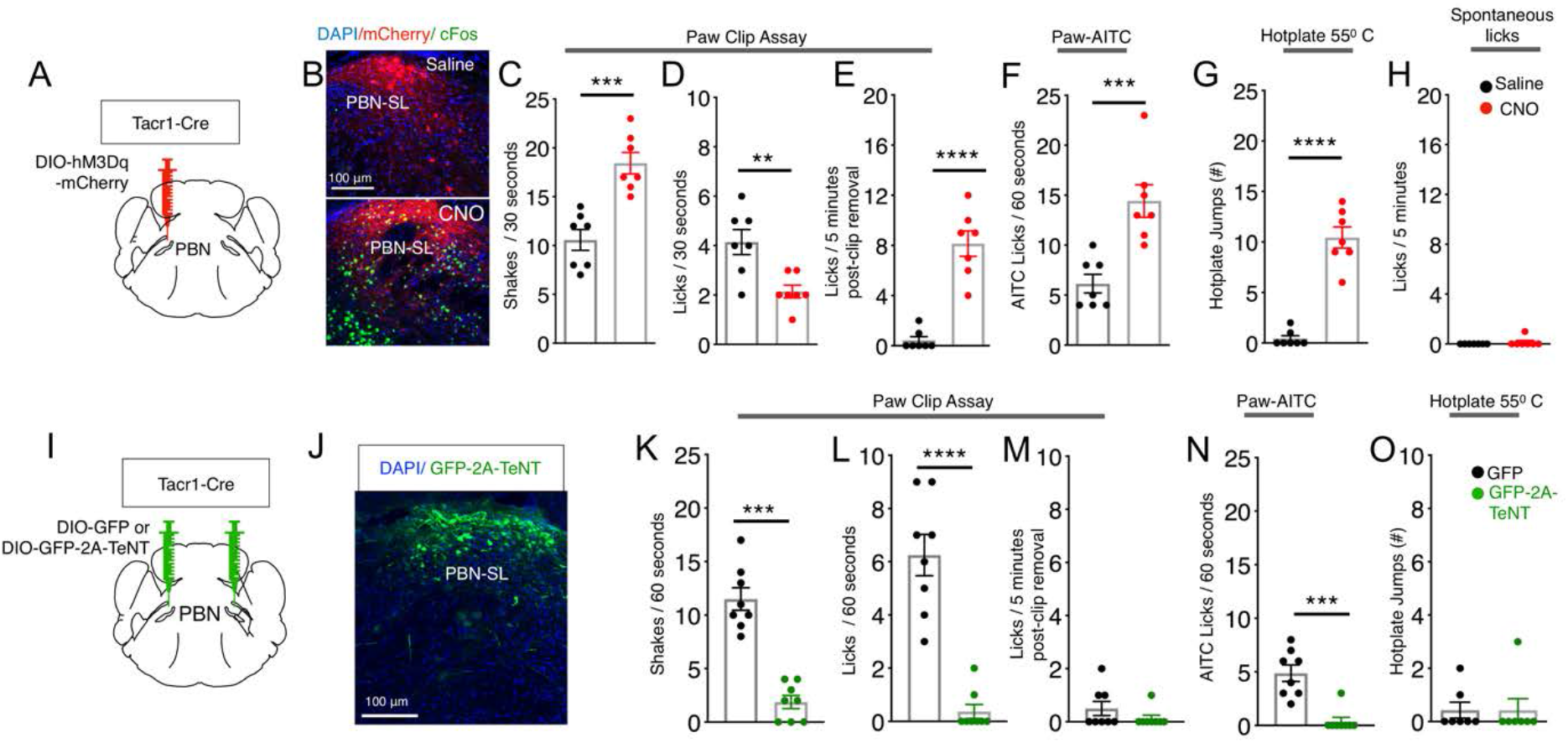
Activation of PBN-SL^Tacr1^ neurons results in a state resembling hyper-vigilance. (A) Chemogenetic activation of PBN-SL^Tacr1^ neurons expressing hM3Dq with CNO (red points) decreased the amount of time mice spent in the center of an open field arena. (B) Mice treated with CNO also display hypersensitivity to gentle punctate touch as measured by a decreased von Frey threshold. CNO treatment did not alter the latency to first response in two thermal assays, Hargreaves (C) or Hotplate (D). n=10 mice. (E-F) Silencing the output from PBN-SL^Tacr1^ neurons with GFP-2A-TeNT (green points) did not alter von Frey thresholds (E) or latency to respond to heat (F). n=8 GFP; n=7 GFP-2A-TeNT. Unpaired two-tailed t test; ns > 0.05, *** p ≤ 0.001, **** p ≤ 0.0001.

**Figure 4 - supplementary Figure 2:**
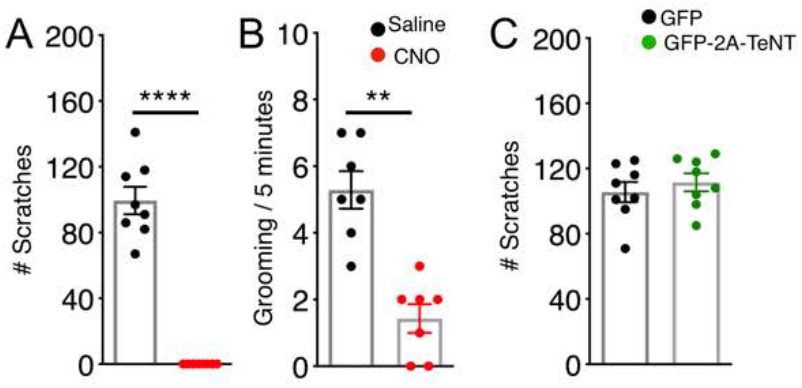
Activation of PBN-SL^Tacr1^ neurons suppresses scratching. (A) Injection of the puritogen chloroquine into the nape of the neck induces robust scratch responses in controls (saline injected mice, black points) that are completely suppressed by chemogenetic activation of PBN-SL^Tacr1^ neurons expressing hM3Dq (CNO, red points). (B) Similarly, activation of these neurons also reduced grooming behavior. n=10 mice (C) Silencing PBN-SL^Tacr1^ neuronal output with GFP-2A-TeNT had no effect on scratching responses. n=8 GFP; n=7 GFP-2A-TeNT. Unpaired two-tailed t test; ns > 0.05, ** p ≤ 0.01, **** p ≤ 0.0001.

